# Sequential ATR and PARP Inhibition Overcomes Acquired DNA Damaging Agent Resistance in Pancreatic Ductal Adenocarcinoma

**DOI:** 10.1101/2024.09.09.609499

**Authors:** Katharine J. Herbert, Rosie Upstill-Goddard, Stephan B. Dreyer, Selma Rebus, Australian Pancreatic Cancer Genome Initiative, Christian Pilarsky, Debabrata Mukhopadhyay, Christopher J. Lord, Genomics Innovation Alliance, Andrew V. Biankin, Fieke E. M. Froeling, David K. Chang

**Affiliations:** Wolfson Wohl Cancer Research Centre, School of Cancer Sciences, University of Glasgow, Glasgow, G61 1QH, UK; West of Scotland Pancreatic Unit, Glasgow Royal Infirmary, Glasgow, G4 0SF, UK; Department of Surgery, Universitätsklinikum Erlangen, Friedrich-Alexander-Universität Erlangen-Nürnberg (FAU), 91054 Erlangen, Germany; Department of Biochemistry and Molecular Biology, Mayo Clinic College of Medicine and Science, Jacksonville, Florida, 32224, USA; CRUK Gene Function Laboratory and Breast Cancer Now Toby Robins Research Centre, The Institute of Cancer Research, London SW3 6JB, UK; Department of Oncology, Beatson West of Scotland Cancer Centre, Glasgow, G12 0YN, UK

**Author notes:** **Corresponding author David K. Chang** Wolfson Wohl Cancer Research Centre, School of Cancer Sciences, University of Glasgow Garscube Estate, Switchback Road, Bearsden, Glasgow Scotland G61 1BD.

**Keywords:** Pancreatic Ductal Adenocarcinoma, acquired treatment resistance, PARP inhibitor, ATR inhibitor, DNA damage response, replication stress, treatment sequence

## Abstract

Pancreatic ductal adenocarcinoma (PDAC) remains the most lethal cancer and will soon be the second most common cause of cancer related death. While regimens containing DNA damaging agents such as FOLFIRINOX and PARP inhibitors have derived clinical benefits for some patients, their efficacy invariably fails over time. This presents a significant clinical challenge, and thus there is an urgent need for novel therapeutic strategies which are able to overcome the acquisition of resistance in PDAC.

Clinically relevant models of treatment resistance were generated from patient-derived cell lines by extended exposure to chemotherapy agents. Synergy scoring, clonogenicity assays, flow cytometry, immunofluorescence and transcriptomic analysis were used to investigate the efficacy of combined ATR and PARP inhibition in re-sensitising resistant PDAC to treatment.

Acquisition of resistance was associated with transcriptomic shifts in cell cycle checkpoint regulation, metabolic control, DNA damage response (DDR), programmed cell death and the replication stress response. Additionally, combined treatment with the ATR inhibitor (ceralasertib), and the PARP inhibitor (olaparib) was synergistic in all models of acquired resistance. Sequential treatment using ceralasertib prior to olaparib was highly effective at low dose for DDR proficient cell lines, whereas DDR deficient models responded better when treated with olaparib first.

We provide *in vitro* evidence of a novel therapeutic strategy to overcome acquired PARP inhibitor and platinum resistance in PDAC by using sequential exposure to ceralasertib and Olaparib. A sequential regimen may be more tolerable and should be investigated clinically to circumvent dose limiting toxicity in concurrent combinations.

## Introduction

Outcomes for pancreatic ductal adenocarcinomas (PDAC) have not changed significantly over the last 50 years with an overall 5-year survival of ∼8%^1^, and is predicted to be the second leading cause of cancer related death by 2030^2^. Reasons for the poor prognosis are multifactorial. Most of the patients present with advanced disease as symptoms are generally non-specific, and many of the patients have rapidly deteriorating performance status, unable to receive any active treatments^3^. In addition, overall response rate for standard chemotherapy is modest and tumours eventually acquire resistance even after a period of treatment response.

Tumour molecular profiling through next generation sequencing has been the foundation of modern oncology to identify targeted therapy that the patient is most likely going to respond to^4,5^. One of the most promising actionable segment are those whose tumours with homologous recombination deficiency (HRD)^6^, which preferentially respond to DNA damaging agents such as platinum and poly (ADP-ribose) polymerase (PARP) inhibitors through synthetic lethality. This approach has significantly improved the outcomes of breast (OlympiAD, OlympiA), prostate (PROfound, PROpel), and ovarian (SOLO-1, PAOLA-1, SOLO-2, and ARIEL3) cancer patients clinically^7–14^. Platinum-based chemotherapy such as FOLFIRINOX has demonstrated clinical efficacy in metastatic^15^ and adjuvant^16^ settings. More recently, maintenance olaparib and rucaparib demonstrated significant progression free survival benefit over placebo in germline *BRCA1/2* carriers and germline and somatic ^17^*BRCA1/2*, *PALB2* PDAC respectively after a period of platinum stabilisation^18,19^. However prolonged exposure to platinum and PARP inhibitors eventually leads to treatment failure over time due to acquired resistance. When acquired resistance happens, unfortunately the prognosis is extremely poor as reported by the RUCAPANC authors^20^, highlighting an urgent unmet clinical need.

Aberrant coordination of events occurring during replication causing replication stress (RS) is a common feature of cancer due to the close relationship of cell cycle and DNA maintenance machinery^21^. An efficient response to and recovery from replication-induced DNA damage is essential to the maintenance of genome wide fidelity, with most cancers developing a degree of enhanced innate tolerance to RS^22^ which in turn is associated with chemotherapy resistance. This may create a dependency on the ataxia telangiectasia mutated and rad3 related kinase (ATR) /CHEK1/WEE1 signalling axis, that coordinates the replication stress response (RSR), and represents an exploitable vulnerability in treatment resistant cancers ^23,24^. We previously reported a RS signature score based on molecular processes involved in maintenance of genomic integrity and demonstrated its correlation to cell cycle checkpoint inhibitor responsiveness such as ATR and CHK in preclinical models of PDAC^25^. Clinically, ATR inhibitors have demonstrated efficacy by differentially targeting tumours with high RS^26^. In addition, early trial results of ATR inhibitor BAY1895344 have shown clinical efficacy in a wide range of heavily pre-treated solid cancers including those acquired resistance to PARP inhibitors^27^. The authors also found evidence of increased DNA damage in on-treatment biopsies, which is unsurprising as acquired PARP inhibitor resistance may be mediated by ATR-induced protection of the replication fork^28^. This further supports the combination of ATR inhibitor to DNA damaging agents as a viable strategy to overcome acquired resistance to DNA damaging agents, even though early phase ATR inhibitor trials showed that intermittent on-off dose scheduling is required to improve tolerability^29^.

Here, we build on our previous work which investigated combination of ATR inhibitor and DNA damaging agents as novel therapeutic strategy to overcome acquired resistance to DNA damaging agents in PDAC, asking whether scheduling ATR inhibition could effectively improve response and circumvent clinical adverse effects.

## Methods

### Chemicals

Chemotherapy agents - cisplatin (Accord Healthcare, London, UK), rucaparib camsylate (Clovis Oncology, Boulder, CO), olaparib/AZD2281 and ceralasertib/AZD6738 (AstraZeneca, Cambridge, UK). Replication stress inducers – hydroxyurea (HU, Sigma-Aldrich), nocodazole (Cayman Chemical, Ann Arbor, MI). All compounds were prepared as stock solutions in DMSO and stored at −80°C until needed. Treatments were serially diluted in PBS immediately before addition to culture media.

### Cell culture

Patient derived cell lines were maintained in high (20%) or low (5%) oxygen conditions at 37°C and 5% CO_2_ as previously described^30–32^. *BRCA2* revertant Capan1 cell lines (described in^33^) was cultured in IMDM with 10% Foetal Bovine Serum (Invitrogen). Cell lines were tested routinely for mycoplasma contamination using MycoAlert PLUS Mycoplasma Detection Kit (Lonza).

### Generation of treatment acquired resistant PDAC cell lines

The TKCC10 patient-derived cell line (parental) was sub-cultured in media containing cisplatin, olaparib or rucaparib over a period of 9 months. Concentration of treatment-containing media was increased until resistant clones were able to survive growth in media containing the 5X the IC_50_ (cisplatin) or 10X the IC_50_ (olaparib or rucaparib) compared to the parent cell line. Where possible, the parent cell line was passage-matched to each resistant clone. Prior to experimental analysis, resistant clones were cultured in fresh treatment-free media for 7-10 days to ensure that results reflected stable change. The *BRCA2* revertant Capan1 cell line (Capan1^BRCA2Revertant^) was previously described in^33^.

### RNASeq expression analysis

RNA extractions were performed using QIAGEN RNeasy Mini kit (Cat#74104) according to manufacturer’s specifications. RNAseq libraries were generated by the Beatson Molecular Technology Unit and sequenced by the Glasgow Precision Oncology Laboratory as described previously^25^. Sequencing quality was assessed with FastQC (v 0.11.9)^34^ and files were processed with fastp (v 0.21.0)^35^ using default settings. Quantification was performed against GRCh38 using Salmon (v 1.4.0)^36^. Salmon quantification results were imported into a DESeqDataSet using DESeq2 (v 1.38.3)^37^. Transcripts were mapped to genes using EnsDb.Hsapiens.v86 (v 2.99.0)^38^. Read count data were filtered to retain only those with normalized counts >= 5 in at least 3 samples (17,403 and 17,080 genes remained in Capan1 samples and TKCC10 samples respectively) and transformed using the DESeq2 ‘vst’ function. All downstream analysis was performed independently for Capan1 cells and TKCC10 cells. Differential expression analysis was performed for the following comparisons: (i) Capan1 parental vs Capan1^BRCA2Revertant^, (ii) TKCC10 parental vs TKCC10 cisplatin, (iii) TKCC10 parental vs TKCC10 olaparib, (iv) TKCC10 parental vs TKCC10 rucaparib. Volcano plots showing significantly differentially expressed (DE) genes (adjusted p-value < 0.05 & absolute(log2FC) > 1) were produced for each comparison with EnhancedVolcano (v 1.16.0)^39^. Heatmaps of DE genes were produced with ComplexHeatmap (v 2.14.0)^40^ and circlize (v 0.4.15)^41^.

### Transcriptome Analysis

Expression-based clustering analysis and heatmaps were generated using Heatmapper (http://www.heatmapper.ca/expression/)^42^. Replication stress scores were calculated for each sample using a previously defined signature of replication stress^25^. Scores were calculated for each gene-set within the replication stress signature with GSVA^43^ and individual scores were summed to obtain an overall replication stress score (RS). Gene Set Enrichment Analysis (GSEA) was performed using Hallmark Gene Ontology (GO) genesets from the Molecular Signatures Database (MSigDB)^44^ with GSEA software version 4.3.2^45^.

### In vitro viability assays

Cells were seeded in 96 well plates and treated after an overnight settle. For sequential treatment experiments, the initial treatment was removed after 24 hr, cells were washed once with PBS, and media containing the second treatment was added to the appropriate wells. Viability was measured 72 hr or 8 days later (depending on the compound) using the CellTiter 96 Aqueous non-radioactive cell proliferation assay (Promega, Madison, WI) with a Tecan Infinite200 Pro plate reader (Tecan Trading AG, Männedorf, Germany). Actinomycin D (Sigma-Aldrich, St Louis, MO), drug vehicle (dimethyl sulfoxide [DMSO]), and media-only controls were performed on each individual plate.

### Colony Formation Assays

Cells were plated as single cell suspensions (500-10,000 cells/well) and left to recover overnight. Treatments were added from concentrated stock solutions directly to seeding media within each well to avoid disturbing the newly attached cells. For sequential treatment, the initial treatment was removed after 24 hrs, wells gently washed with PBS, and the final treatment added to the appropriate well. In the case of experiments using cisplatin, cells were exposed for 24 hr, after which treatment containing media was removed and replaced with fresh media. Colonies were grown for 14-21 days, stained with crystal violet and counted using an automated GelCount plate scanner (Oxford Optronix, Banbury, UK). The plating efficiency (PE) [PE = Average Colony Number/Cells plated] and the surviving fraction (SF) was calculated using [SF = PE_Treated_ /PE_Untreated_]. The significant difference between nonlinear modelled curves was calculated by two-way ANOVA, with survival as the dependent variable and treatment conditions as the independent variables.

### Immunoblotting

Cell lysates were prepared in RIPA lysis buffer (Thermo Scientific) with protease inhibitors (Roche) and phosphatase inhibitors (Sigma) and protein concentration was determined using the BCA assay (Thermo Scientific). Proteins were separated by SDS-PAGE and transferred to PVDF membranes, which were probed by overnight incubation at 4 °C in primary antibody solution. Targets were detected via HRP-conjugated secondary antibodies exposed to chemiluminescence reagent (Millipore) and imaged using a Licor Odyssey XF Imager (Li-Cor, Lincoln, Nebraska). Antibodies used were anti-SLFN11 (ab121731, Abcam); anti-ATR (ab10312, Abcam); anti-Rad51C (NB100-177, Novus Biologicals); anti-Cyclin E (sc-247, Santa Cruz); anti-Rad51 (ab133534, Abcam); anti-pRPA(S4/S8) (ab243866, Abcam); anti-Actin (CST#3700, Cell Signalling Technology); HRP linked anti-rabbit and anti-mouse IgG (CST#7074 & 7076, Cell Signalling Technology). Images were processed with Image Studio Analysis software v 4.0 (Li-Cor, Lincoln, Nebraska).

### Replication fork stall recovery

Cells were treated with hydroxyurea overnight to induce fork stalling. Treatment media was removed, cells were washed twice in prewarmed PBS and released into media containing 100 ng/mL nocodazole to prevent re-entry into G1. Samples were obtained every 2 hours for 10 hrs following release and processed immediately for cytometric analysis. For each HU-treated sample, a non-HU-treated control sample was prepared identically to standardise for potential interference by nocodazole.

### Cell cycle analysis

Samples were labelled by incubating cells with BrdU (Sigma) at 20 μM in growth medium for 30 min and fixed in ice-old 70% ethanol at 4°C. S phase immunostaining was performed after pepsin/acid digestion using anti-BrdU (1:100, Beckton Dickson) primary antibody, and AlexaFluor488 conjugated secondary antibody (1:500, Molecular Probes). Following incubation, stained cells were pelleted and incubated in a solution containing propidium iodide (Sigma Aldrich) stain (20 μg/mL) and RNase A (200 μg/mL) for 30 min. Samples were scanned using an Attune NxT cytometer (Invitrogen) at a flow rate of 150-300 events/sec.

### Foci counting

Cells were grown on coverslips at sub confluent densities and treatments initiated during logarithmic growth phases. At the relevant time points, nonchromatin bound nuclear protein was removed by preincubation of coverslips with CSK extraction buffer (100mM NaCl, 300mM Sucrose, 3 mM MgCl2, 10 mM PIPES (pH 7.0), 1 mM EGTA, 0.5% Triton X100) for 10 min on ice, after which samples were fixed with 4% paraformaldehyde for 15 min. Coverslips were blocked with 2% goat serum before overnight incubation with primary antibody at 4°C [anti-Rad51 (1:500, Santa Cruz), anti-53BP1 (1:1000, Cell Signalling Technology), anti-phospho-RPA2(Ser4/8)(1:5000, Abcam), and anti-phospho-Histone H2A.X (Ser139) (1:2000, Millipore)], followed by the corresponding fluorescent secondary antibodies [AlexaFluor555 (1:1,000, Molecular Probes) or AlexaFluor488 (1:1,000, Molecular Probes)] for 1 h at room temperature. Nuclei were counterstained with 0.5μg/ml 4′,6-diamidino-2-phenylindole (DAPI; Sigma-Aldrich), and coverslips were mounted using Vectashield (Vector Laboratories). Z stack images were randomly acquired under identical parameters with a Zeiss LSM780 confocal microscope (Zeiss) using a ×63 oil immersion objective. Data are represented as mean ± SEM of two independent experiments evaluating at least 200 nuclei in each experiment.

### Statistical analysis

Zero-interaction potency (ZIP) synergy scoring was calculated using SynergyFinder 3.0^46^. For 53BP1, pRPA2, γH2AX and RAD51 foci evaluation, images were analysed using FIJI (v2.9.0)^47^. Statistical analysis and graphs were produced using GraphPad Prism v9.4.1 (GraphPad Software, San Diego CA). Foci counting experiments were performed twice, with >4 randomly assigned fields captured by confocal microscopy per condition. For colony formation assays, data are representative of three independent experiments from triplicate wells. Data were analysed by two-way ANOVA, with Bonferroni, Tukey’s, or Sidak’s post-tests used for multiple comparisons. Spearman’s rank correlation tests were used to assess homoscedasticity, and checks were performed for Gaussian distribution during analysis. Differences were considered significant at a P value of <0.05, and unless stated, all results are presented as mean ± SD or mean ± SEM as indicated.

## Results

### Higher expression of RS response genes is associated with poor survival in PDAC

We and others have previously shown that high RS correlated to poorer prognosis in PDAC patients^25^ (Supplementary Figure S1A). To further investigate the prognostic effects of RSR genes, we used expression of genes from the MSigDB Reactome Pathways R-HSA-176187, R-HSA-5656169, and R-HSA-73894 (activation of ATR activation in response to replication stress, termination of translation synthesis, DNA repair), on the ICGC PACA-AU cohort (Fig 1A). Tumours with higher expression of *SLFN11, RAD51C, CDK2, LIG3* and *POLA1* were associated with better prognosis (Fig 1B; Supplementary Fig S1B), whereas tumours with increased expression of RSR genes such as *Cyclin E1/2, CHEK1/2, RAD51, ATR, DCLRE1/ARTEMIS* and *RPA3* alone or in combination had worse prognosis, making them ideal novel therapeutic targets.

**Figure 1:**
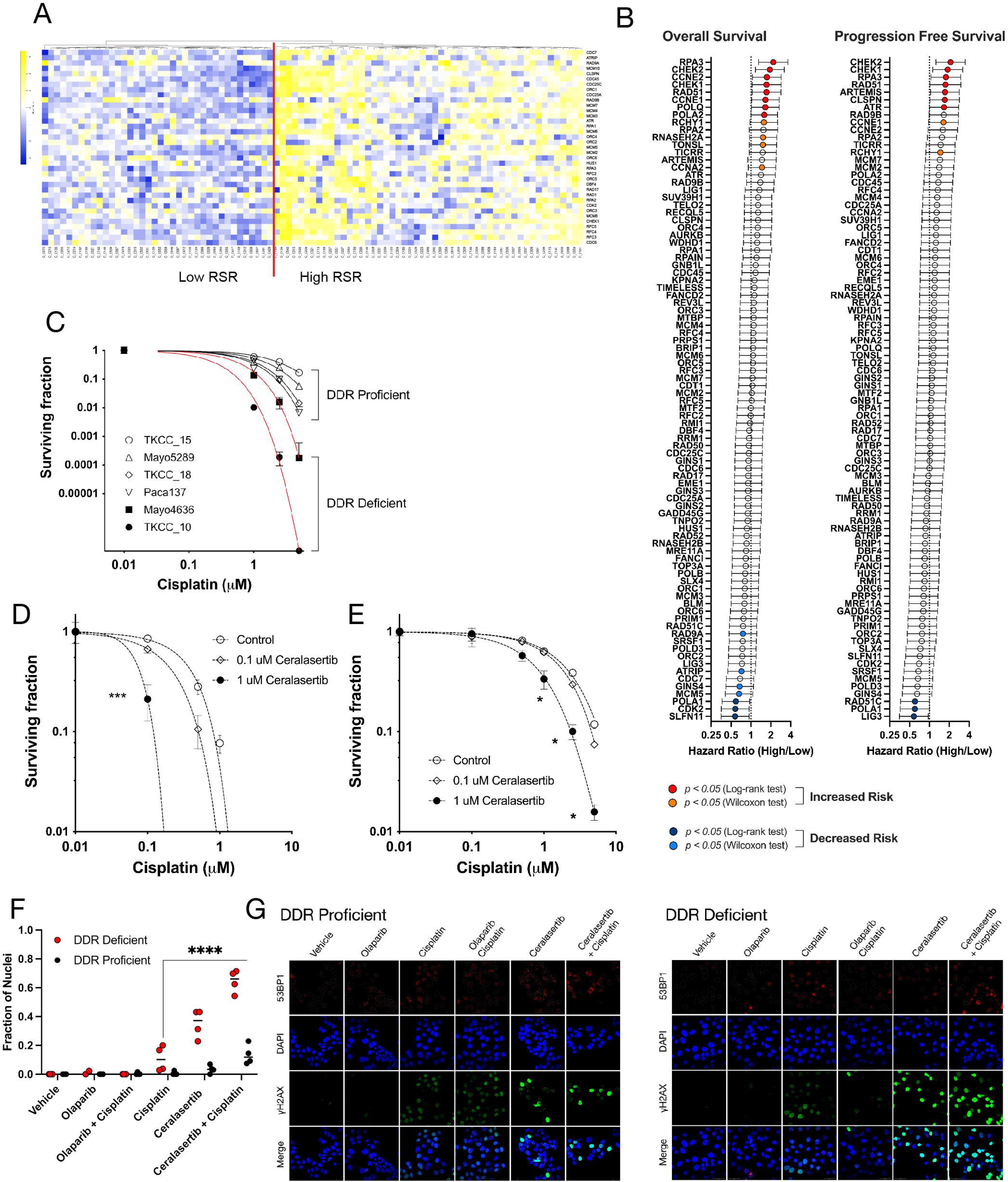
RS and DDR vulnerabilities are prognostic in pancreatic ductal adenocarcinoma. **A.** Gene expression and survival data in primary pancreatic ductal adenocarcinomas (PDAC) from ICGC data sets were interrogated to uncover prognostic relationships between replication stress response (RSR) gene expression and DNA damage response (DDR) proficiency. Manhattan clustering of transcriptomic data was used to classify tumours as having high or low RSR potential. **B:** Log-Rank testing of survival data was performed to calculate overall and disease-free survival hazard ratios for RSR genes. Survival ratio (± 95% CI) for higher relative expression is shown; genes with lower risk are indicated by blue symbols, and those having significantly increased risk are in red. **C:** Clonogenicity assays were performed on cisplatin treated PDCLs, and surviving fraction plotted relative to the plating efficiency of the untreated sample. Curves from DNA damage proficient and DNA resistant cell lines are indicated. Ceralasertib and cisplatin treatment combinations were tested on cisplatin sensitive (**D**) and cisplatin resistant (**E**) TKCC10 patient derived cell lines using colony formation assays. Nonlinear regression analysis was performed to determine the effect of overnight treatment for each condition, and individual datasets analysed by 2-way ANOVA with Tukey’s test for multiple comparisons (*, *p* < 0.05; ***, *p* < 0.001). **F and G:** DDR deficient (TKCC10) and DDR proficient (TKCC26) were exposed to 1 μM concentrations of olaparib, cisplatin, or ceralasertib for 24 hr, probed for γH2AX and imaged by confocal microscopy. Fraction of pan-nuclear γH2AX stained nuclei were analysed by 2-way ANOVA with Tukey’s test for multiple comparisons (****, *p* < 0.0001). Graph is representative of 2 independent experiments, with quantification performed on 4 confocal images containing approximately 20 nuclei per region of interest.

### Inhibition of Replication Stress response enhances sensitivity to Cisplatin in PDAC

We previously showed that RS score correlates to ATR inhibitor responsiveness^25^ (Supplementary Fig S1C) and concluded that both RS and ATR signalling capacity determines tumour viability. To further explore therapeutic strategies, we treated a panel of patient-derived cell lines (PDCLs) that have been comprehensively characterised at the genomic and transcriptomic level, using cisplatin in combination with an ATR inhibitor, ceralasertib. We demonstrated preferential cisplatin response in DDR deficient models (Fig1C). The addition of overnight treatment with ceralasertib further increases sensitivity to cisplatin in clonogenicity assays regardless of DDR status (Fig1D, 1E), and was associated with a significant increase in pan-γH2AX staining, consistent with replication catastrophe due to combination treatment (Fig 1F, 1G). Based on these results, we further explored if manipulation of RSR with ATR inhibition can overcome acquired treatment resistance in PDAC.

### Generation and Characterisation of Acquired Resistance preclinical models of PDAC

Development of acquired resistance to platinum and PARP inhibitor treatment is common^48^ and is associated with poor prognosis. We modelled acquired cisplatin and PARP inhibitor resistance in PDCLs using prolonged exposure to treatment, or *BRCA2* reversion mutation through CRISPR-mediated reversion of gene function.

#### Prolonged Exposure

We sub cultured TKCC10 (a DDR deficient *BRCA1* mutated PDCL with relatively high basal RS, based on prior analysis^25^) in treatment-spiked growth media (cisplatin, olaparib or rucaparib) at increasing concentrations over a 9-month time period to produce resistant colonies which were able to tolerate a 5-10 fold higher concentration of the agent compared to parental (Fig 2A-C). We then performed RNASeq and analysis for each of the acquired resistant cell lines (Fig 2E-H; Supplementary Fig S2A, SC-E). This demonstrated down regulation of genes implicated in programmed cell death (*CKMT1A*, *PTPN13*) (Fig 2I) and upregulation of genes promote cell migration (*NTN1*, *JCAD*, *AMOTL1*, *ATP10D*) (Fig 2J) that are common in all acquired resistant models. While olaparib and rucaparib acquired resistant models generated largely unique transcriptomic profiles, they shared ∼ 20% of the differentially expressed genes. These included genes that increase multidrug resistance (*TCEAL9*↑), membrane trafficking (*EPN3*↑), cell growth and proliferation (*LIPG*, *BBF2H7*), and promotion of an immune repressive microenvironment (*MFAP5*↓, *CSF1*↑), and downregulation in those associated with redox homeostasis (*PRXL2A*↓), and DNA damage response pathway control (*ALDH1A1*↓) (Fig 2I & J) explaining some of the underlying acquired resistance mechanism.

**Figure 2.**
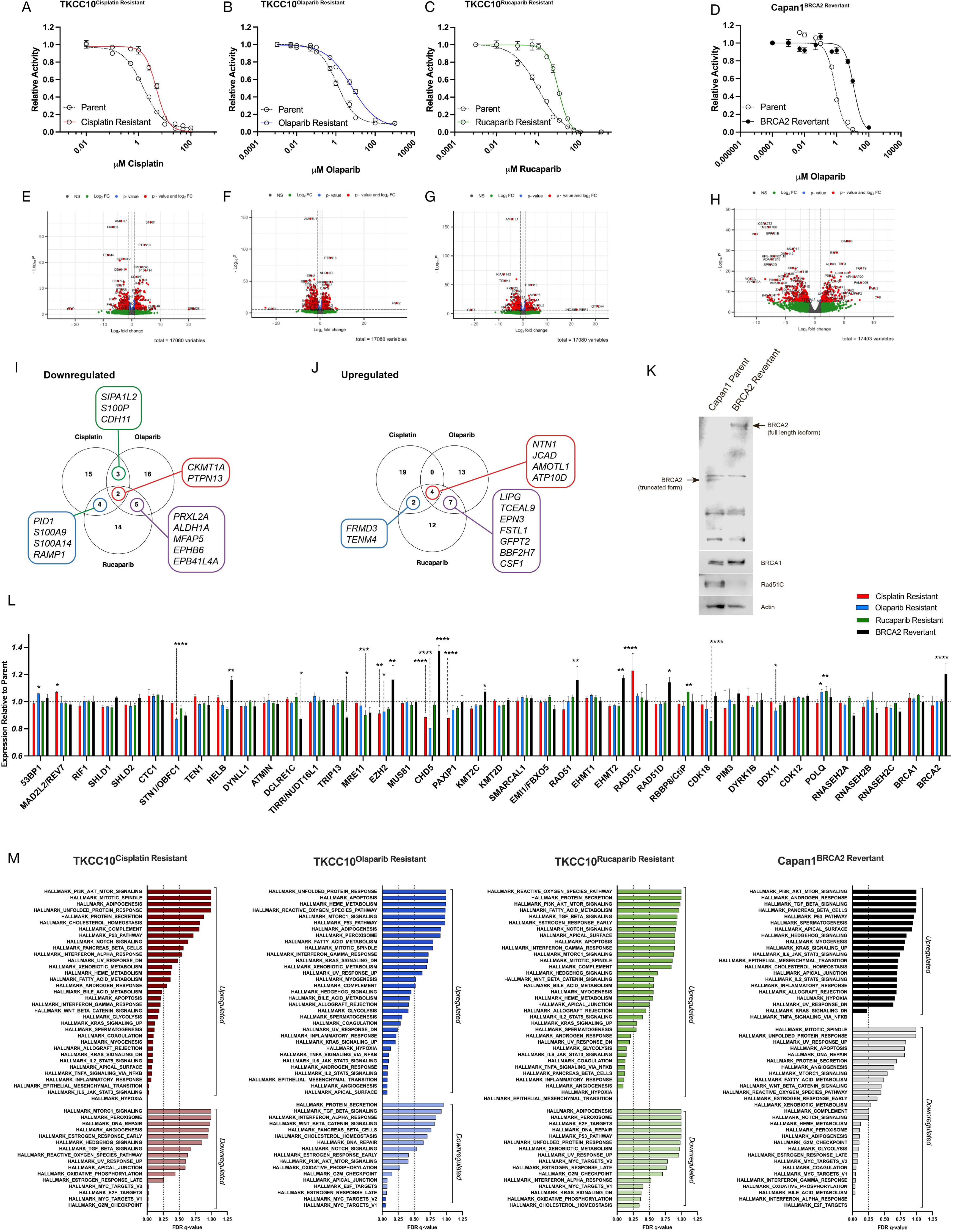
Modelling acquired resistance in PDAC cell lines. **A:** Dose-response assay results for cisplatin resistant (Red); **B:** Olaparib resistant (Blue); **C:** Rucaparib resistant (Green); and **D:** *BRCA2* Revertant (Black). Cell viability was determined using MTS assay and calculated relative to vehicle control. Curves from parent cell lines are indicated by dashed lines (Uncoloured dots). Panels show representative results from 3 independent experiments. Relative activity for cisplatin assay was recorded after 72 hours, and after 8 days’ exposure for olaparib and rucaparib. RNASeq was performed on TKCC10 Parental and Treatment Resistant cell lines; and Capan1 Parental and BRCA2 Revertant cell lines (*n*=3). Volcano plots are of differentially expressed genes from cisplatin resistant (Panel E), olaparib resistant (Panel F), rucaparib resistant (Panel G), and BRCA2 Revertant (Panel H) cell lines compared to passage-matched Parental samples. Downregulated (Panel I) and upregulated (Panel J) differentially expressed genes for TKCC10 resistance cell lines compared to the parental cell line. Panel K: Immunoblots of Capan1 Parent and BRCA2 Revertant cell lysates probed for BRCA2, BRCA1, RAD51C and Actin protein expression. Multiple BRCA2 isoforms are displayed, arrows indicate truncated and full-length isoforms. Panel L: Difference in expression of resistance-associated genes relative to parental cell line was analysed by 2-way ANOVA with Dunnett’s test for multiple comparisons (cisplatin resistant, red; olaparib resistant, blue; rucaparib resistant, green; BRCA2 Revertant, black). Results are from 3 independent experiments (*, p < 0.05; **, p < 0.01; ****, p < 0.0001). Panel M: Enrichment analysis was performed on treatment resistant cell lines using Hallmark gene sets.

#### Mutation Reversion

Reactivation of gene function by reversion mutation is a significant cause of acquired resistance to DNA damaging agents in breast, ovarian and pancreatic cancer^49–51^. Capan1 is a commonly used PDAC cell line harbouring a *BRCA2* frameshift mutation causing truncation of BRCA2 at its DNA binding domain leading to homologous recombination repair deficiency and impedes recovery from replication stress-induced lesions^52^. To recapitulate a reversion mutation seen clinically, we used a Capan1^BRCA2Revertant^ cell line, which harbours a CRISPR-engineered reversion mutation in *BRCA2* that partially restores gene function^33^, which in turn results in a 10-fold lower sensitivity to olaparib (Fig 2D, Fig 2K). We then performed RNASeq on the Capan1^BRCA2Revertant^ cell line and revealed significant increase in expression of genes involved in growth and proliferation (*ROBO1*↑), immunosuppression (*HHLA2*↑), and epithelial to mesenchymal transformation (*ADAM8*↑*, FAM3B*↑), and decrease in genes involved in membrane trafficking (*FAM109B*↓), programmed cell death (*CALHM2*↓), cell migration (*IQGAP2*↓) (Fig 2H; Supplementary S2F) when compared to parental cell line which only carries a dysfunctional *BRCA2* allele. Previous studies in HR deficient ovarian and breast cancers that have acquired resistance to platinum chemotherapy and/or PARP inhibitor ^53,54^ have implicated those genes enriched at collapsed replication forks during ATR inhibition^55^. Our analysis showed changes in 20 DDR and RS associated genes, with 3 affected genes (*EZH2*, *CHD5*, and *POLQ*) common to more than one resistance type. (Fig 2L).

All acquired resistant cell lines had reduced RS scores compared to parental (Supplementary Fig S2G), and hallmark pathways analysis revealed downregulation of the G2/M checkpoint, and E2F and MYC targets, and upregulation of hypoxia pathways, promotion of epithelial-mesenchymal transition, and IL2, IL6, TNFα, and KRAS signalling in acquired resistance cell lines (Fig 2M).

### Acquired resistance is associated with impaired RS tolerance and recovery

We next investigated how acquired resistant PDCLs respond to and recover from exogenous RS induced by treatment with hydroxyurea (HU), a ribonucleotide reductase inhibitor. Cell lines were treated overnight with 1mM HU to induce replication fork stalling at the G1/S border, and replication progression was tracked by analysing the BrdU positive population in early S (S1), mid-S phase (S2), or late S (S3) following removal of the inhibitor (Fig 3A).

**Figure 3.**
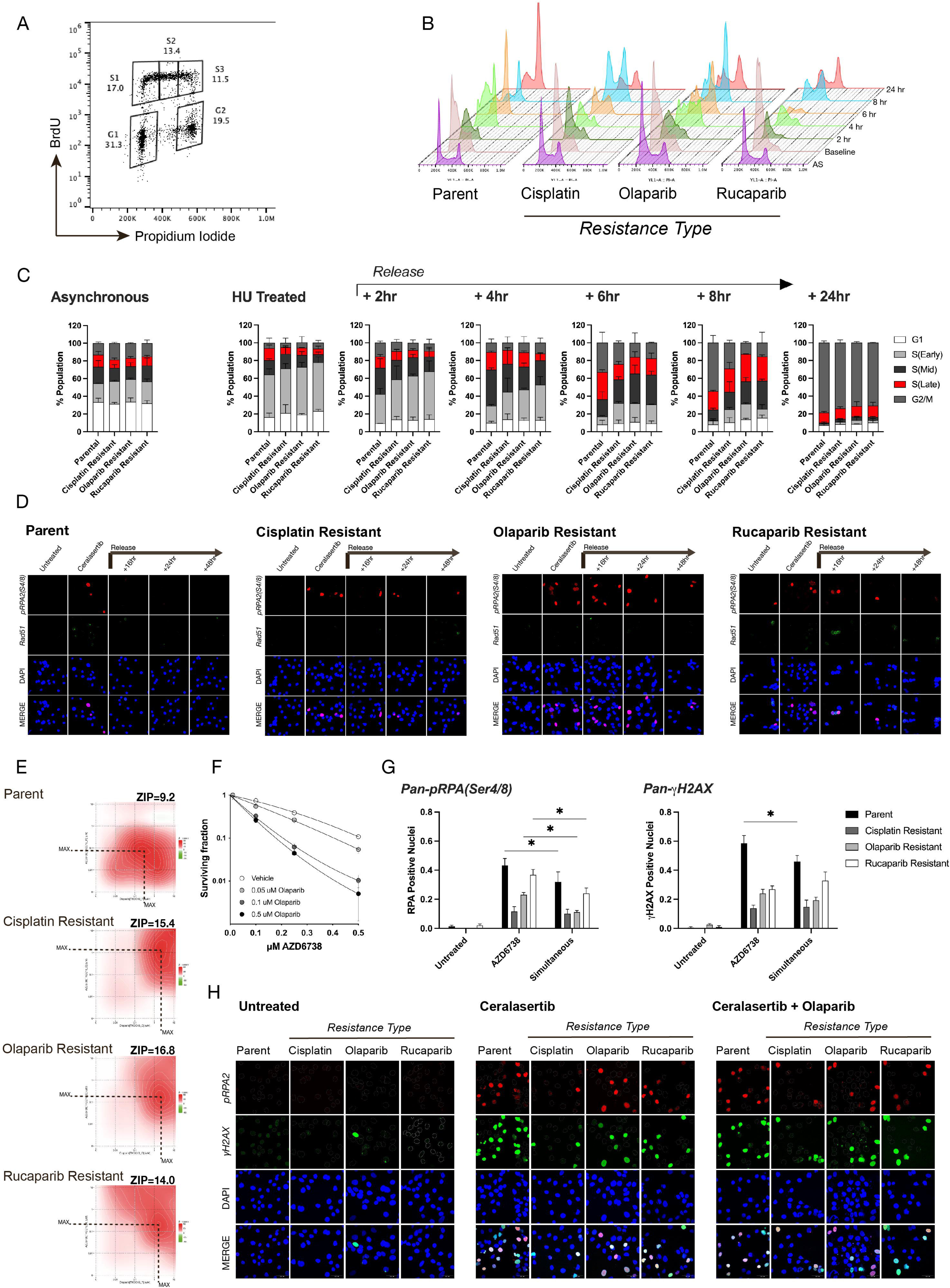
AZD6738 and Olaparib are synergistic in treatment resistant PDAC. **A:** TKCC10 parent and treatment resistant cell lines were synchronised in G1/S by overnight treatment with hydroxyurea and replicating cells were tracked at 2 hourly intervals following release using BrdU uptake. Cell cycle was assayed by propidium iodide staining (**B**). Progression of replicating cells through S phase was analysed in 3 phases – Early S(S1), Mid S(S2), and Late S(S3); gating strategy is indicated. Percentage of the total population at each cell cycle phase is shown for each timepoint (**C**). TKCC10 parental and resistant cell lines were treated with ceralasertib and olaparib in a 5 × 5 matrix of dose combinations (10-fold dilutions, 0-10 μM). Cell viability was assayed by MTS after 8 days’ exposure. Drug synergy was calculated using the interaction potency (ZIP) model across the dose matrix (**E**). Clonogenicity analysis of the effect of ceralasertib and olaparib in combination was performed on the TKCC10 cell line using treatment across a 0-0.5 μM dose range. Surviving fraction was calculated relative to the plating efficiency and analysed by 2-way ANOVA. Survival curves were generated using the linear quadratic model (**F**). The response of resistant cell lines to ceralasertib and olaparib combination treatment and ceralasertib monotherapy (1 μM concentrations for each, 24 hr exposure) was assayed using Pan-nuclear γH2AX and Pan-nuclear pRPA2 staining as markers for replication catastrophe (Panels G & H). Results were analysed by 2-way ANOVA with Tukey’s test for multiple comparisons (*, *p* < 0.05; *ns*, non-significant; results from 2 independent experiments).

Although we found each acquired resistant cell line had the same percentage of cells in each cell cycle phase as the parental cell line at rest (Fig 3A), S phase progression is significantly altered in acquired resistance models. Parental TKCC10 cells enter S phase 30 minutes following release (S1), with most of the population transitioning through mid-S (S2) at approximately 4 hours post-release and reaching late S (S3) in 6-8 hours. In acquired resistance PDCLs, the transition through S phase is much slower at 8 hours post-release, with ∼ 20% of cells failing to fully complete S phase and transition into G2 (Fig 3B & C) demonstrating impaired tolerance and recover from exogenous RS. Considering the importance of ATR/CHEK1 axis activity in RSR, we treated acquired resistant models with ceralasertib for 72 hours and showed decreases sensitivity on viability assay in line with a lower RS score (Supplementary Figure S3A&B). To test the differences in recovery potential with shorter exposure, we treated these cell lines with 1μM of ceralasertib for 5 hours.

While we saw the same initial amount of replication-associated damage in the form of pan-nuclear pRPA2 between the parental and the acquired resistant cell lines (Fig 3D), the parental cell lines recovered faster after the inhibitor was washed out, with pRPA2 foci retained for at least 48 hours in all of the resistant cell lines, and RAD51 foci significantly increased in rucaparib acquired resistance model (Fig 3D; Supplementary Fig S3C). These indicate RSR triggered by ATR inhibition is retained following washing out the agent, which opens the possibilities of concurrent or sequential combination with other agents.

### ATR and PARP inhibition are synergistic

Considering the reduction in endogenous RS as shown by reduced RS signature score and reduced sensitivity to ATR inhibition in acquired resistance models, we next asked if manipulation of the RS response by combining ceralasertib and olaparib can re-sensitise and overcome acquired resistance. Olaparib is known to activate the G2/M checkpoint, causing an increase in the G2 population of exposed cells^56^. To assess the effects of olaparib on cell cycle, we treated a panel of PDAC PDCLs with single agent olaparib, and this slowed the production of replication associated damage in acquired resistant cell lines by stalling cell cycle progression at the G2/M border regardless of DDR status (Supplementary Fig S3D), making it ideal to combine with ATR inhibitor.

We then use low dose olaparib and ceralasertib simultaneously over 10-14 day and found significantly reduced clonogenicity for the parent cell line at sub-micromolar concentrations (Fig 3F). The combination was synergistic for both parental and acquired resistant cell lines (Fig 3E) even though the parent cell line remained more sensitive than resistant cell lines (MAX Synergy scores: Parent = 9.179; Cisplatin resistant = 15.378; Olaparib resistant = 14.02; Rucaparib resistant = 16.775; Fig 3E). We also observed an increase in the G2 population, albeit for cell lines with high RS scores, activation of the S phase checkpoint by ceralasertib appears to dominate (Supplementary Fig S3D). In all but the cisplatin resistant model, overnight exposure to combination treatment reduced replication associated DNA damage (in the form of RAD51 and pRPA foci) when compared to ceralasertib monotherapy (Supplementary Fig S3C). Using pan-nuclear γH2AX staining as an indicator for replication catastrophe we found that simultaneous combination treatment did not significantly increase replication stress-associated cell death in resistant cell lines when compared to ceralasertib alone (Fig 3G, Fig 3H), and produced a small but significant reduction in replication catastrophe measured for the parent cell line.

Considering that sensitivity to olaparib by ceralasertib depends on a combination of S phase exit and replication stress signature, we next investigated if sequentially scheduling treatment can influence sensitivity.

### Scheduled ATR inhibition enhances PARP inhibitor sensitivity in resistant PDAC

We first assessed clonogenicity on the Parental TKCC10 cell line by treating with one agent followed by treatment with the second agent with each treatment completely washed out with PBS after overnight exposure. We found treatment sequence order significantly influenced the outcome. When ceralasertib was used first, there was no dose-dependent effect on clonogenic synergy, whereas sensitisation increased when cells were treated with olaparib first (Fig 4A). We then assessed the effect of sequential treatment on acquired resistance models with 24hr exposure to either ceralasertib or olaparib prior to treatment with the alternative agent using a viability matrix across the same dose range as Fig 3E.

**Figure 4.**
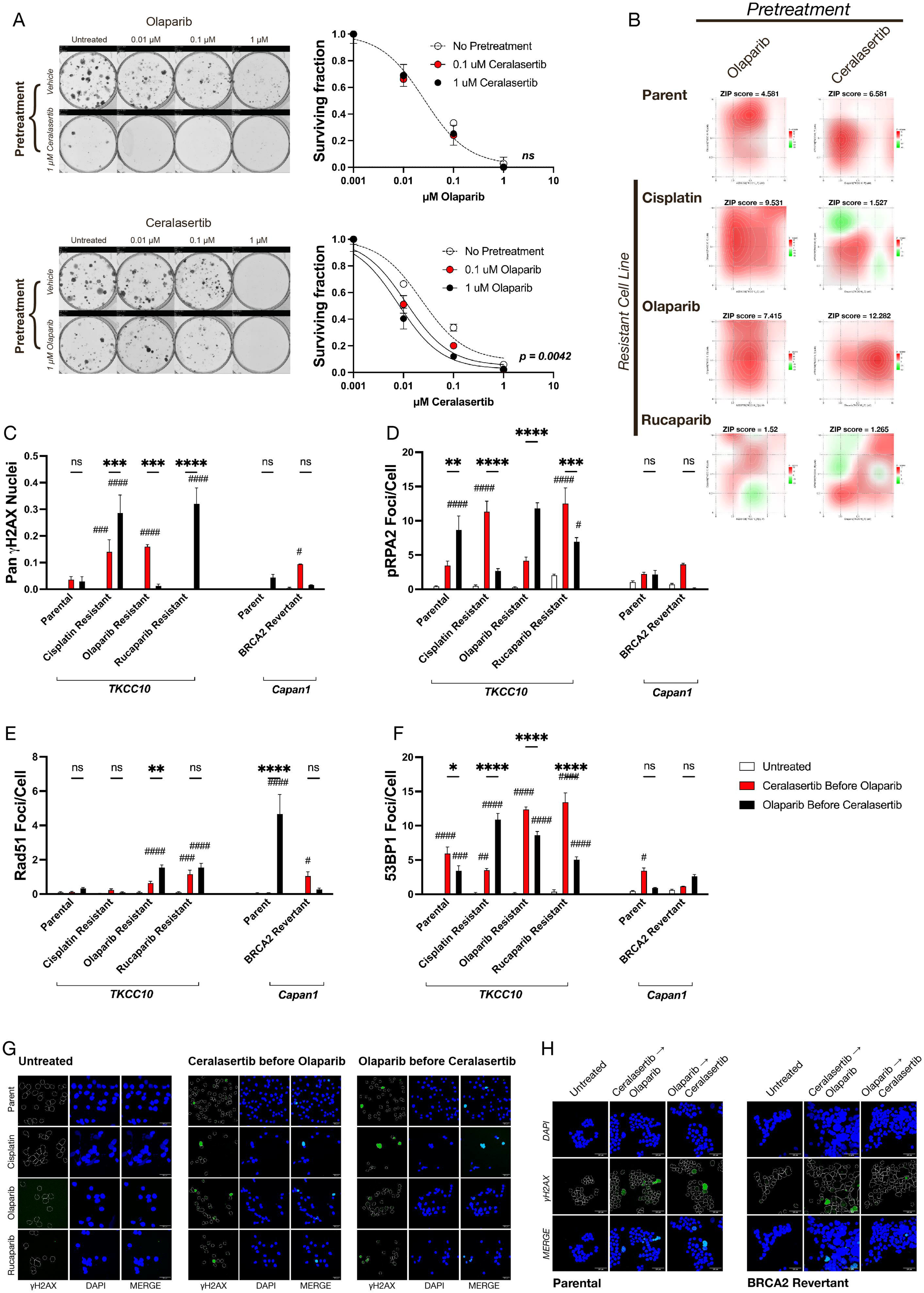
Ceralasertib sensitises acquired resistant PDAC to PARP inhibition. **A:** Clonogenicity assay results from the TKCC10 cell line after sequential ceralasertib and olaparib treatment using a matrix of sequential dose combinations. Results were analysed by nonlinear regression relative to the surviving fraction of the vehicle for pretreated (red symbol) or non-pretreated (clear symbol) conditions. Graphs are representative of 2 independent experiments and p values for significantly different curves are indicated. Representative clonogenicity assay images are shown. **B:** TKCC10 (Parent) and Cisplatin, Olaparib or Rucaparib resistant cell lines were sequentially treated with ceralasertib and olaparib in a 5 × 5 matrix of dose combinations (10-fold dilutions, 0-10 μM) in either order. Cell viability was assayed, and drug synergy calculated using the interaction potency (ZIP) model. Dashed lines indicate maximum synergy. TKCC10 acquired resistance cell lines and Capan1 Parental and BRCA2 Revertant cell lines were seeded at sub-confluent densities on coverslips and treated for 24 hours with 0.1 μM concentrations of ceralasertib or olaparib; or sequential treatment with each treatment in either order. Bars indicate percentage of cells for each condition with Pan-γH2AX stained nuclei (**C**), pRPA2 foci per cell (**D**), Rad51 foci per cell (**E**), and 53BP1 foci per cell (**F**). The effect of combination was analysed by 2-way ANOVA with Tukey’s test for multiple comparisons. Conditions were compared to vehicle control (p < 0.001, ###; p < 0.0001, ####), and effect of alternative treatment order was compared (*, p < 0.05; **, p < 0.01; ****, p < 0.0001). Results are from 2 independent experiments, with at least 4 different fields of view captured for image analysis. Representative confocal images from Pan-γH2AX staining are shown in TKCC10 acquired resistance cell lines (**G**) and Capan1 Parental and BRCA2 Revertant (**H**).

The efficacy of ceralasertib was enhanced 10-fold by prior exposure to olaparib using the acquired cisplatin resistant model. With the acquired olaparib resistant model, sensitivity to olaparib was enhanced by previous ceralasertib treatment, and with the acquired rucaparib model sensitivity to rucaparib increased with prior exposure to ceralasertib (Fig 4B; Supplementary Fig S4A and S4B). In either case the opposite sequential order did not produce a synergistic response of the same magnitude. These data demonstrate that sensitivity to each PARP inhibiting agent is dependent on adaptation to prior exposure and indicate that the element driving resistance should be used as backbone in combination with ATR inhibition.

We then investigated the underlying mechanisms contributing to different synergy and sensitivity of various drug sequence and regimens in different models of acquired resistance. The percentage of pan-nuclear γH2AX positive cells generated when ceralasertib was used prior to olaparib was significantly greater for the olaparib resistant model, however in cisplatin and rucaparib acquired resistance models, the opposite was true (Fig 4C, Fig 4G). This pattern of enhanced DNA damage sensitivity associated with specific treatment order is in line with the response for each resistance model seen in Figure 4B. To better understand if sensitivity to treatment order is connected to each model’s capacity to recover from replication-associated DNA damage, we next measured the expression of nuclear markers for single strand breaks (pRPA2 foci), and double strand breaks (53BP1 foci) as well as Rad51. Expression of these markers are in-line with the treatment sequence and regimens producing peak synergy for each acquired resistance models (Fig 4D-F). In addition, for both Olaparib and Rucaparib acquired resistance models, significantly more Rad51 foci were observed with sequential treatment, indicating a greater reliance on these factors for SSB and DSB stabilisation and repair.

We next investigated if the same patterns of treatment sensitivity are seen in models of resistance acquired through reversion mutation mechanism using Capan1 and Capan1*^BRCA2revertant^* isogenic cell lines. Inhibiting the RSR in the DDR proficient Capan1*^BRCA2revertant^*cell line with ceralasertib prior to olaparib increased residual DNA damage and restored sensitivity to olaparib in viability assay, produced 3-fold higher synergy (Supplementary Figure S4C), and more replication catastrophe (Fig 4C-D, Fig 4H) than by using olaparib first in the sequence. Foci counting assays confirms these findings, showing increased DDR activity with ceralasertib before olaparib for the Capan1*^BRCA2revertant^*cell line (Fig 4E-F). Taken together, this demonstrates that treatment sensitivity can potentially be enhanced by combined manipulation of the DDR and RSR pathways in a sequential manner, and is effective regardless of the mechanism driving resistance.

### Sequential ATR and PARP inhibition is effective in PDAC PDCLs

We next investigated if sequential ceralasertib and olaparib treatment could be effective in the endogenous resistance setting by further screening a panel of 10 PDCLs with varying degree of DDR status and RS signature as previously described^25^ for synergy and relative clonogenicity. In these models, *BRCA1* and *BRCA2* loss through mutation are the main cause of DDR deficiency (Fig 5A), and SLFN11 expression positively correlated to RS signature scores (Fig 5B, Supplementary Figure S5A). We found olaparib first then ceralasertib achieved significantly higher synergy scores in DDR deficient / RS high models using viability assays. However, in DDR proficient / RS low models, ceralasertib first then olaparib was more synergistic (Fig 5C; Supplementary Fig S5B). Results for clonogenicity mirrored the outcome of synergy scoring assays, with olaparib first sensitising DDR deficient PDCLs to ceralasertib, and ceralasertib first enhancing sensitivity to olaparib for DDR proficient PDCLs (Fig 5D). However, the role of RS signature is less evident in the different treatment sequence, as cell lines with lower RS scores typically have poor plating efficiency, and colony formation capacity in this assay format. Taken together, the different sequential treatment regimen can potentially be effective in other settings including DDR proficient and a range of RS score models.

**Figure 5.**
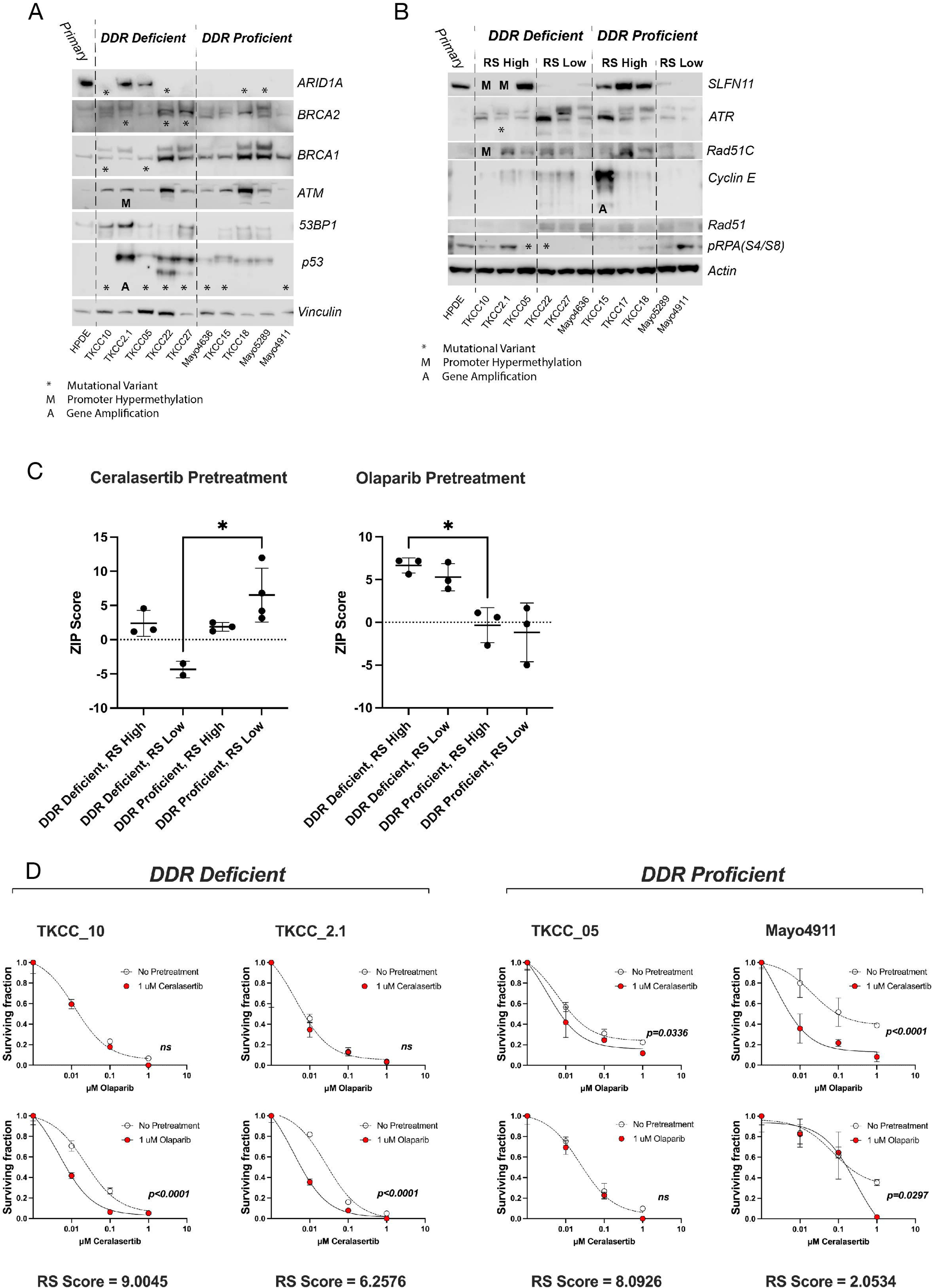
Sequential ATR and PARP inhibitor treatment is effective in PDAC cell lines. **A:** Immunoblot from patient-derived cell lines probed for endogenous expression of DNA damage response markers. Cell lines are grouped according to DNA damage response (DDR) proficiency. Mutational variants (*), promoter hypermethylation (M) and amplification (A) of relevant genes for individual cell lines are indicated. **B:** Immunoblot from patient-derived cell lines probed for replication stress response markers. Cell lines are grouped according to DNA damage response (DDR) proficiency and replication stress (RS) score (High or Low). Mutational variants (*), promoter hypermethylation (M) and amplification (A) of relevant genes for individual cell lines are indicated. **C:** ZIP synergy score analysis for patient-derived cell lines after sequential treatment with ceralasertib and olaparib in a 5 × 5 matrix of dose combinations (10-fold dilutions, 0-10 μM). Results are grouped by DNA damage response proficiency and replication stress score (high, H; or low, L), and were analysed by 2-way ANOVA with Sidak’s multiple comparison test (* p < 0.05). **D:** Graphical representation of clonogenicity assay results from sequential ceralasertib and olaparib-treated patient-derived cell lines. Results were analysed by nonlinear regression relative to the surviving fraction of the vehicle for pretreated (red symbol) or non-pretreated (clear symbol) conditions. Graphs are representative of 2 independent experiments and p values for significantly different curves are indicated.

## Discussion

DNA damaging agent containing regimens such as FOLFIRINOX and NALIRIFOX^57^ as well as PARP inhibitors such as olaparib and rucaparib have demonstrated survival benefits in patients with PDAC, and in some cases exceptional responses that translated to prolonged survival. However, patients who derived meaningful benefits remains the minority, and inevitably, even the responders eventually acquire treatment resistance, which is associated with poor prognosis. Significant efforts have been generated by the scientific and clinical communities to overcome acquired resistance, either with combinatorial regimens, or targeting other aspects of the DNA damage response pathways. Some of the limitations to deploy combinatorial strategy has been dose limiting toxicity as demonstrated by the VIOLETTE clinical trial, where the olaparib plus adavosertib (AZD1775, WEE1 inhibitor) arm was terminated early due to toxicity^58^. The same trial also terminated the concurrent olaparib plus ceralasertib arm due to no additional benefit with the addition of ATR inhibitor over PARP inhibitor alone in acquired platinum resistance triple negative breast cancer^58^.

Similarly, while the CAPRI trial using the same olaparib plus ceralasertib regimen as VIOLETTE demonstrated encouraging evidence of clinical benefit in PARP inhibitor acquired resistant high grade serous ovarian cancer^59^, there was some tolerability challenges with concurrent administration of the two investigational drugs, likely indicating the requirement of reducing dose intensity compared to respective monotherapies, or intermittent on-off treatments.

Here, we provide *in vitro* evidence of novel therapeutic strategy of sequential ceralasertib than olaparib to overcome acquired PARP inhibitor and platinum resistance in PDAC models by exploitation of RSR in terms of tolerance and recovery as therapeutic vulnerability. We further demonstrated the sequence of DNA damaging agents and ceralasertib matters as well as the context of the prior line DNA damaging agent exposure. Finally, we showed that we were able to use this sequential treatment regimen of ceralasertib first then olaparib to sensitise a large panel of PDAC models to olaparib regardless of their DNA damage response status and / or innate replication stress level. While current clinical investigations have focused around concurrent doing of Olaparib and ceralsertib with variable dose intensity and on-off regimens, the sequential regimen we have presented here may be a viable alternative strategy to circumvent the challenges of dose limiting toxicity in the combinatorial regimens trailed clinically and should be tested further in well-designed clinical trials for patients who acquired resistance to DNA damaging agents.

