## Supplementary Information for "Sequential ATR and PARP Inhibition Overcomes Acquired DNA Damaging Agent Resistance in Pancreatic Ductal Adenocarcinoma"

**Supplementary Figure Legends**

Supplementary Figure S1.

Heatmaps showing Manhattan clustering of ICGC bulk tumour (Panel A) into high or low expression of gene ontology (GO) pathways involved in the DNA damage response and cell cycle control. Delineation of tumour samples into high and low replication stress (RS) score are shown.

Panel B: A high or low replication stress recovery score was calculated for the ICGC cohort using the combined Z score from genes identified as high or low risk in Figure 1B. Survival analysis was performed for overall survival and progression free survival using the median value as a cutoff. Curves were compared by Logrank Test (p values shown).

IC50 values for 72hr exposure to the ATR inhibitor ceralasertib plotted against the replication stress signature scores using cell lines derived from various PDAC tumours (Panel C). Linear regression line is shown.

Supplementary Figure S2.

Panel A: Principal Component Analysis (PCA) map of RNASeq samples from parental cell line (TKCC10_P), cisplatin resistant cell line (TKCC10_C), olaparib resistant cell line (TKCC10_O), and rucaparib resistant cell line (TKCC10_R); and Panel B Principal Component Analysis (PCA) map of Capan1 parental (Control) and Capan1 BRCA2 Revertant (Treatment) cell lines. Points refer to 3 independently generated replicates.

Differential gene expression (Top 50) heatmaps for TKCC10 cisplatin resistant (Panel C; TKCC10_C), olaparib resistant (Panel D; TKCC10_O), and rucaparib resistant (Panel E; TKCC10_R) cell lines relative to vehicle treated parental control (TKCC10_P).

Panel F: Differential gene expression (Top 50) heatmap for Capan1 BRCA2 Revertant (Capan1_BRCA2_Rev) relative to the Parental cell line (Capan1_P).

Panel G: Graphical representation of replication stress scores (RS) calculated for TKCC10 parental and resistant cell lines, and Capan1 Parental and BRCA2 Revertant cell lines (Bars indicate mean ±SEM; n = 3).

Supplementary Figure S3.

Panels A & B: Dose-response viability assay curves for 72 hours ceralasertib exposure in TKCC10 acquired resistance (Panel A) and Capan1 BRCA2 Revertant (Panel B) cell lines.

Panel C: TKCC10 parent and treatment resistant cell lines were seeded on coverslips and treated for 5 hours with Vehicle or 1 μM ceralasertib. Treated cells were released from treatments by washing the coverslips twice with PBS. Culture media was replaced and treated coverslips were sampled at 16, 24, and 48 hour timepoints following release. Coverslips were probed for Rad51 foci and pan-nuclear phospho-RPA(S4/S8) expression. Nuclei were counterstained with DAPI. Coverslips were mounted and imaged by confocal microscopy (5 images/condition sampled randomly, representative images shown).

Panel F: Cell cycle cytometric analysis of cell lines with high replication stress score (TKCC02.1); and low replication stress score (TKCC26) after overnight treatment with 1 μM concentrations of olaparib, ceralasertib, or both olaparib and ceralasertib combined. Nuclear DNA content was stained with propidium iodide (PI), and cell cycle populations were analysed using FlowJo software. Images are representative of 2 replicates.

Supplementary Figure S4.

Graphical representation of the zero-interaction potency (ZIP) scores for TKCC10 (Parent) and Cisplatin, Olaparib or Rucaparib resistant cell lines sequentially treated with ceralasertib and olaparib (Panel A); TKCC10 acquired resistance cell lines treated with ceralasertib and rucaparib (Panel B); and Capan1 Parent and BRCA2 Revertant cell lines treated with ceralasertib and olaparib (Panel C).Treatments were administered in either order, with an initial 24 hr pretreatment, followed by 8 days exposure to the Post-treatment in a 5 $\times$ 5 matrix of dose combinations (10-fold dilutions, 0-10 μM).

Supplementary Figure S5.

Panel A: Heatmap showing high and low clustering of replication stress (RS) scoring data from ICGC patient derived cell lines using gene sets involved in the DNA damage response and cell cycle control.

Panel B: Combined synergy scores (CSS) for patient-derived cell lines after sequential treatment with ceralasertib. Uncoloured points represent values from cell lines pretreated with olaparib, and red point indicate values from those pretreated with ceralasertib. Results are grouped by DNA damage response proficiency and replication stress score (high, H; or low, L), and were analysed by 2-way ANOVA with Sidak's multiple comparison test (* $p < 0.05$; ** $p < 0.01$; *** $p < 0.001$).

### Supplementary Figure S1

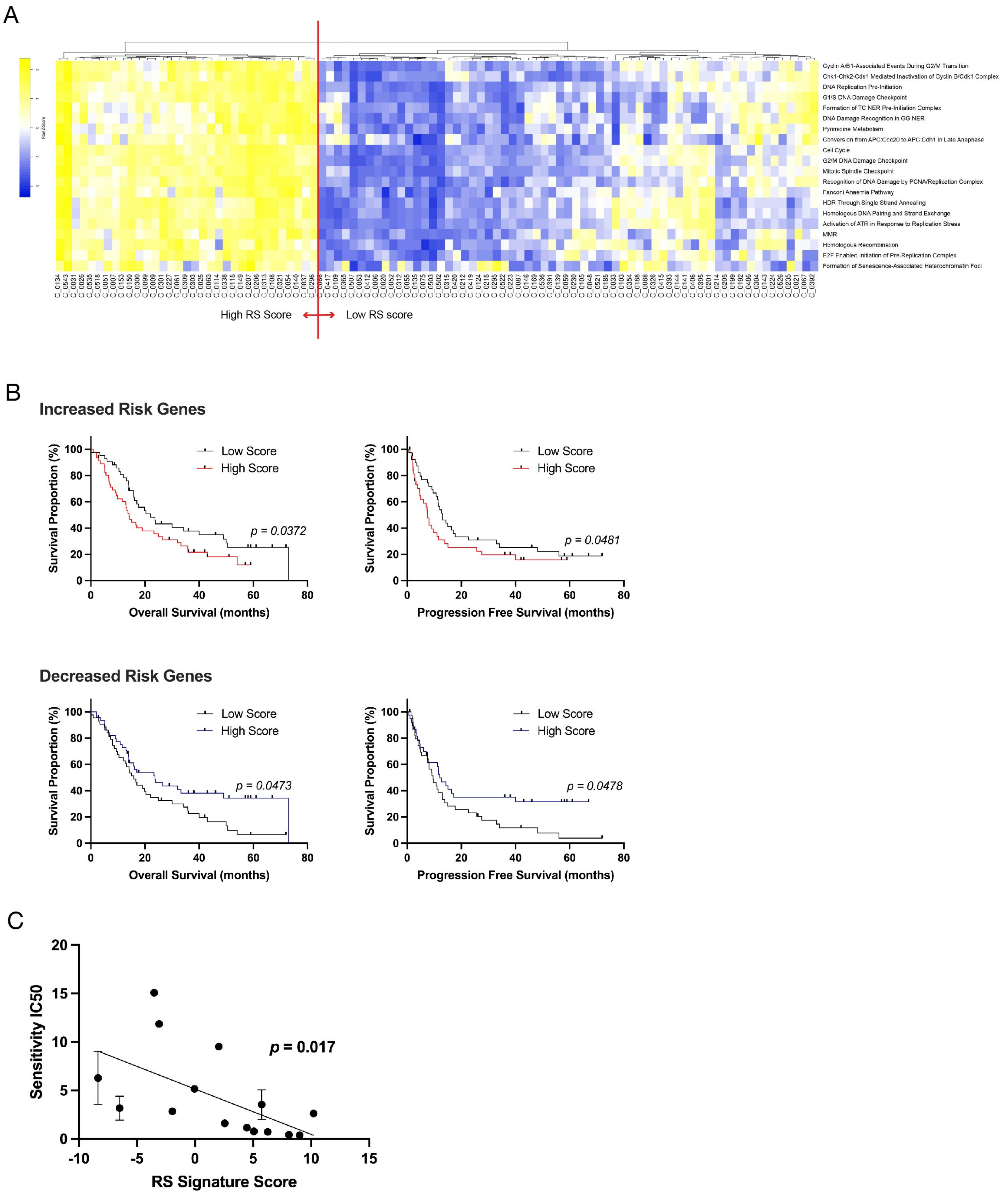

### Supplementary Figure S2

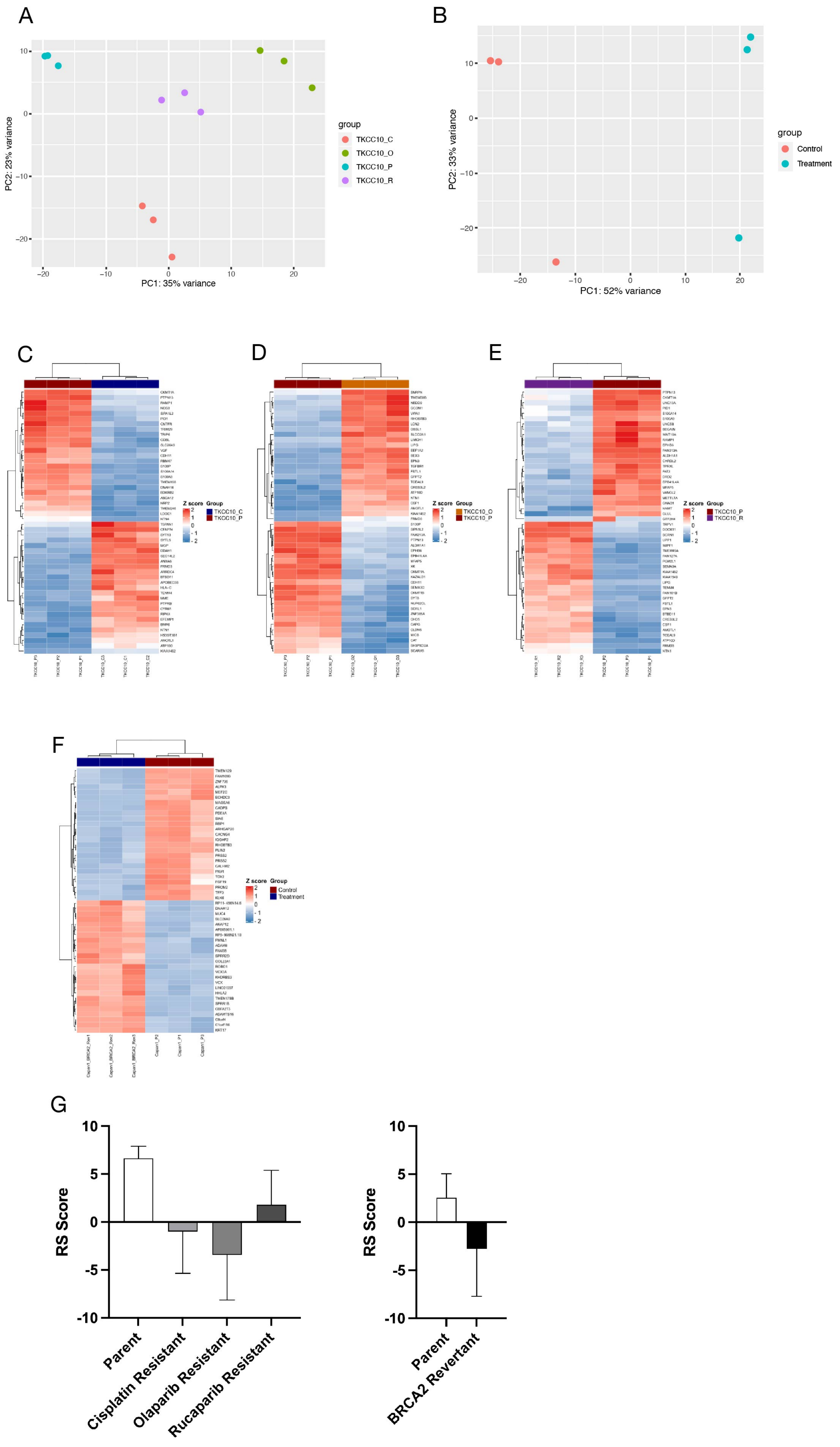

### Supplementary Figure S3

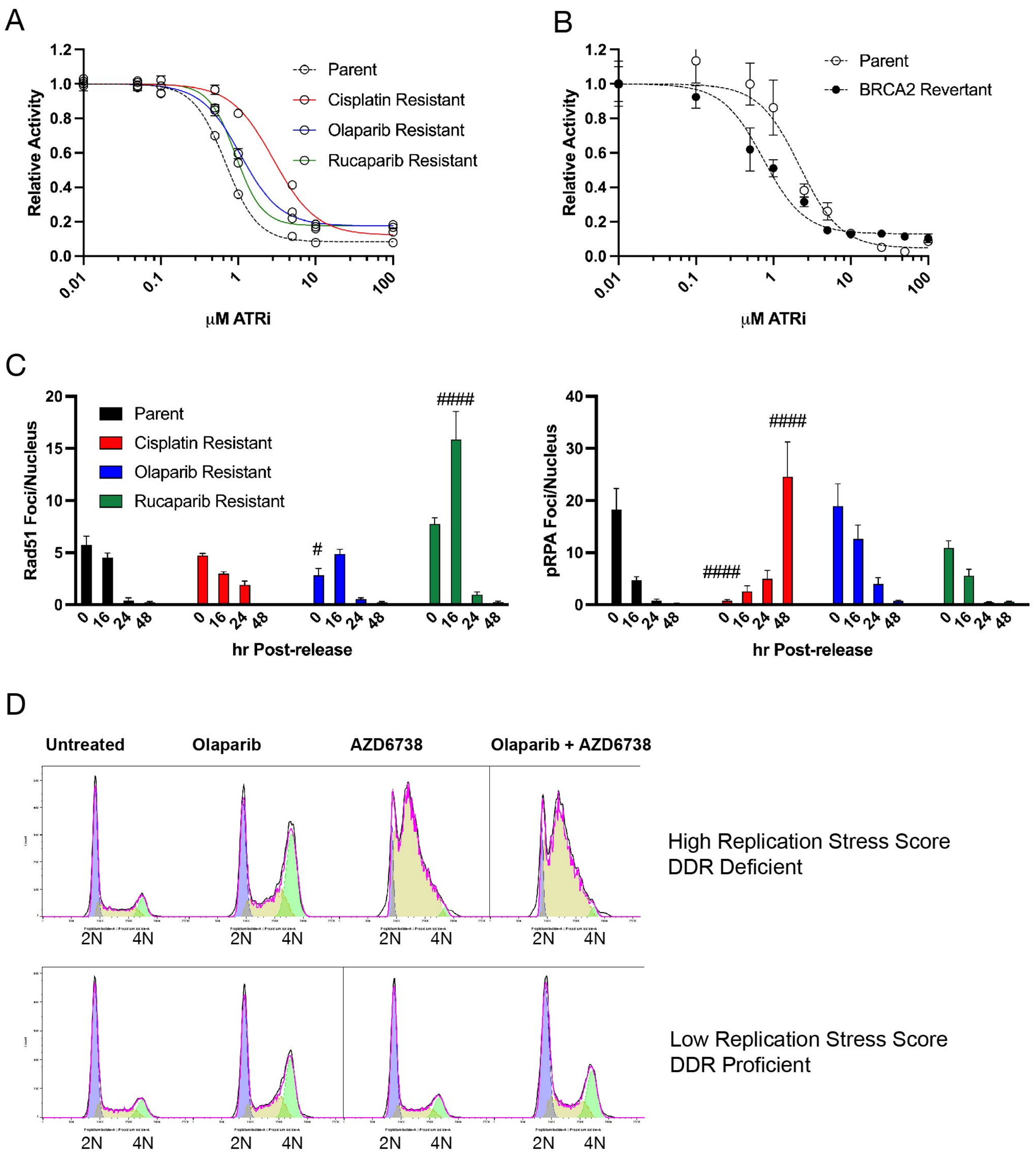

Supplementary Figure S4

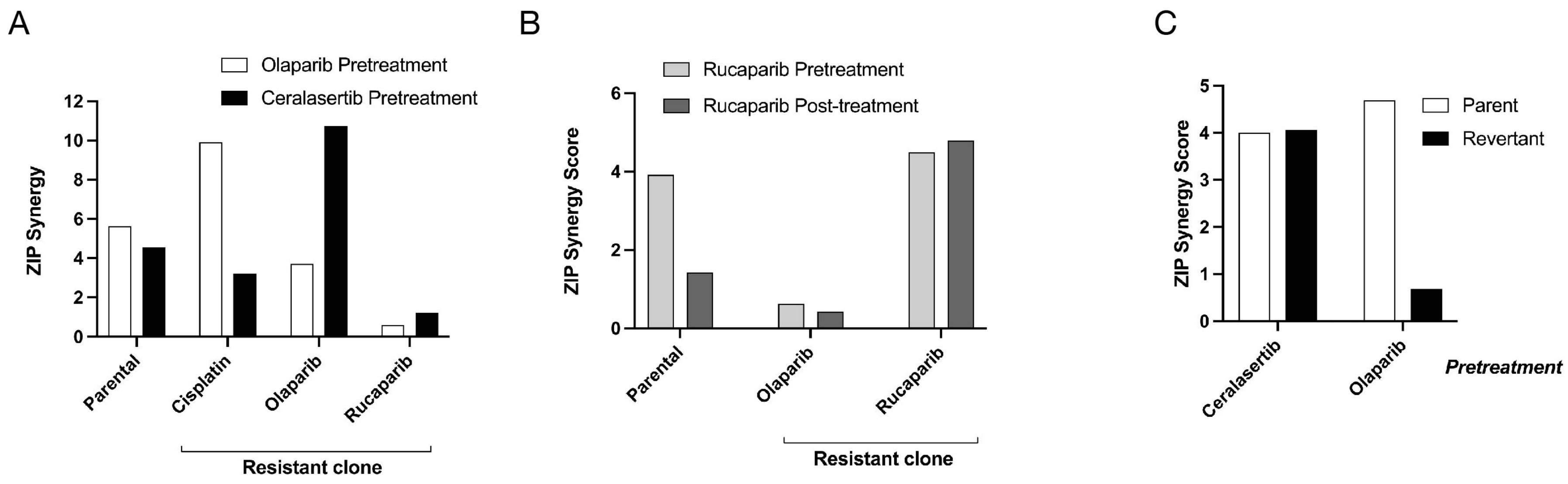

### Supplementary Figure S5

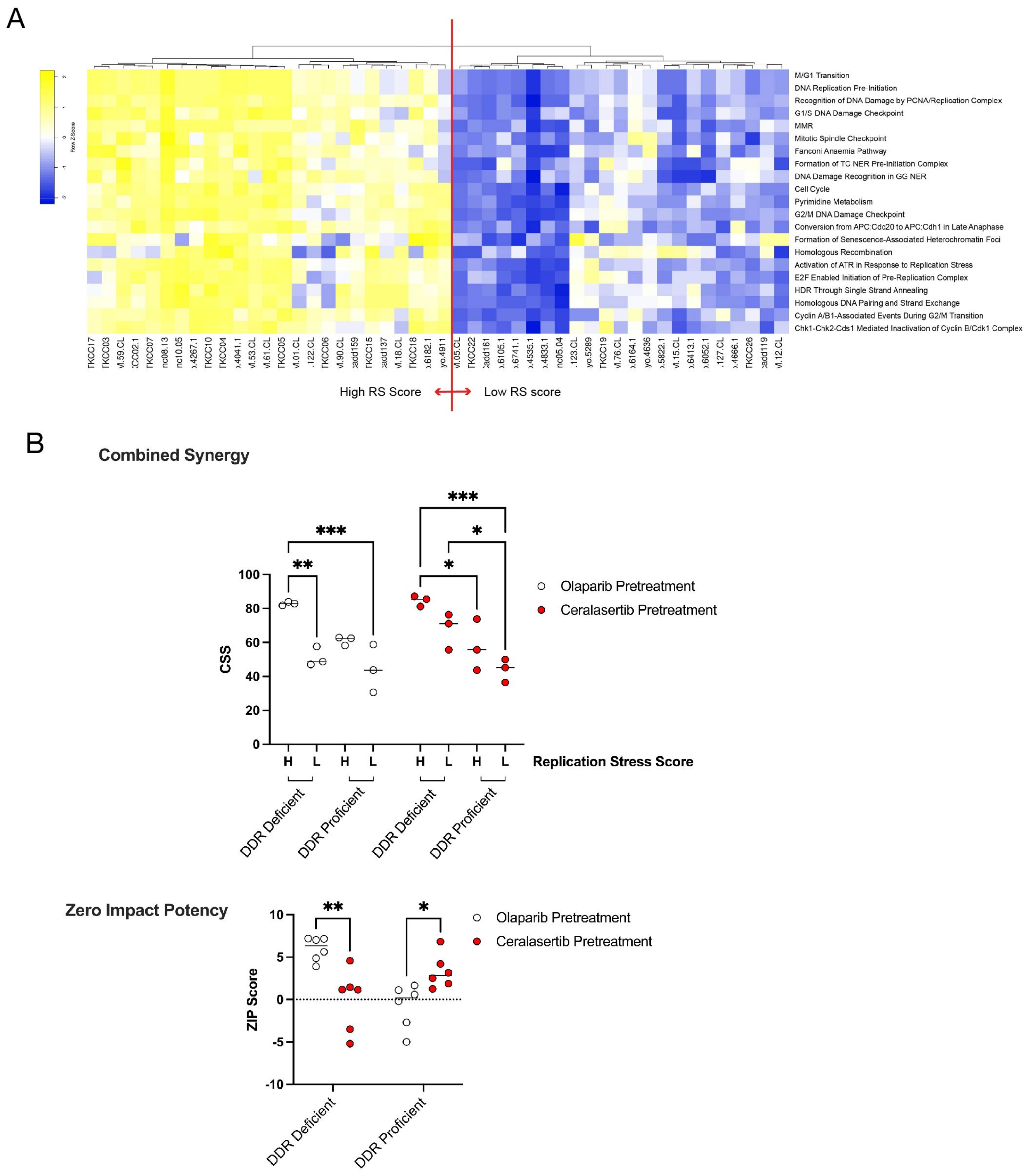
